# BRD1 haploinsufficiency alters early neuronal programming and disrupts maturation in human induced glutamatergic neurons

**DOI:** 10.64898/2025.12.19.695380

**Authors:** Julie G. Donskov, PJ Michael Deans, Dimitrios Pediotidis-Maniatis, Jacob E. Høgfeldt, Anders D. Børglum, Mark Denham, Kristen J. Brennand, Per Qvist

**Author notes:** Corresponding author: Dr. Per Qvist, Ph.D. Department of Biomedicine, Aarhus University Wilhelm Meyers Allé 4, building 1242 8000 Aarhus, Denmark. These authors share last authorship (Kristen J. Brennand, Per Qvist).

## Abstract

BRD1 is an epigenetic regulator implicated in neurodevelopmental and psychiatric disorders, yet its role in human neuronal differentiation, maturation, and function remains poorly understood. Here we show that *BRD1* haploinsufficiency disrupts early neuronal programming, resulting in accelerated maturation and altered neurodevelopmental trajectories in human induced glutamatergic neurons. Transcriptomic profiling reveals an early shift toward neuronal identity, characterized by downregulation of pluripotency markers and persistent upregulation of genes involved in synapse assembly and organization, including *GRIA3*. Despite this, *BRD1*^+/-^ neurons form significantly smaller synapses and display increased neuronal activity. Our findings highlight BRD1 as a key regulator of neurodevelopmental timing and synaptic maturation, and network activity reinforcing growing evidence that disruptions in chromatin-mediated control of differentiation and synaptic organization contribute to neurodevelopmental disorders.

## Introduction

Disruptions in brain development and function contribute to a spectrum of disorders with overlapping genetic and neurobiological underpinnings. Schizophrenia (SZ), bipolar disorder (BPD), and autism spectrum disorder (ASD) are traditionally categorized as distinct conditions, yet growing evidence suggests they share common developmental vulnerabilities (1). This overlap is supported by genomic studies, which reveal significant genetic correlation and shared polygenic risk across major neuropsychiatric (NPD) and neurodevelopmental (NDD) disorders (1,2). These findings underscore the etiological complexity of these conditions, with risk variants spanning multiple biological pathways (3–7).

While genetic variants directly affecting synaptic function likely influence specific aspects of neural circuitry (4,7–12), others contribute to regulatory processes that more broadly support neuronal maturation, network formation and functional circuits in the brain(13–16). These developmental processes rely on precisely coordinated molecular mechanisms, including chromatin remodeling and epigenetic regulation. Mutations in genes coding for epigenetic modifiers and chromatin remodelers are overrepresented in neuropsychiatric disorders, and disruptions to such regulatory systems are increasingly recognized as key contributors to the clinical heterogeneity and pleiotropic effects observed across neuropsychiatric conditions (17,18).

*BRD1* (Bromodomain Containing 1) encodes an epigenetic regulator that plays a pivotal role in chromatin recruitment to modulate histone acetylation (19). It has emerged as a compelling candidate gene governing spatio-temporal transcriptional regulation in the developing brain. Genetic variations in the *BRD1* gene have consistently been associated with SZ and BPD (20–26), while rare disruptive mutations have been identified in SZ and ASD patients (4,23,27–29). Beyond direct associations, the *BRD1* interactome is enriched with genes implicated in neurodevelopment and mental health (30). Emphasizing the importance of BRD1-regulated transcription in normal brain development and function, *BRD1* has been associated with cortical surface area in the general population (31,32), and studies in rodent and porcine preclinical knockout models have linked *Brd1* haploinsufficiency with neuro-morphometric changes, alterations in brain cell composition, and cognitive and behavioral changes with broad translational relevance to psychiatric disorders (33–38).

While preclinical knockout models have contributed significantly to our understanding of the role of *BRD1* in psychopathology (33–39), their clinical translatability remains limited.

In this study, we investigate the molecular and functional impact of *BRD1* haploinsufficiency on neuronal development. Using human pluripotent stem cell (hPSC)-derived glutamatergic neurons (iGLUTs), we assess the consequence of BRD1 deficiency across key stages of progenitor transition, differentiation, cell-type specification, migration and maturation. By integrating transcriptomic and functional analyses, this study aims to elucidate the role of BRD1 in neurodevelopmental processes and its potential contribution to the pathophysiology of neuropsychiatric disorders.

## Materials and methods

### Study design

Commercial H9 human pluripotent stem cells (hPSCs) were genetically modified using CRISPR/Cas9 to generate a heterozygous *BRD1^+/-^* hPSC line. *BRD1^+/-^* and WT hPSCs were then converted into mature neurons via neurogenin-2 (hNGN2)-induction and kept in culture for up to 42 days. Cultures were characterized using bulk RNA sequencing on D7, D21 and D28, and hallmark features of neuronal development were assessed using immunocytochemistry (ICC) followed by high-content imaging microscopy on D7, D14 and D21. Finally, as a readout of neuronal function, neural activity was repeatedly measured from D21-D42 using a multielectrode array (MEA). A schematic presentation of the study design are shown in **Figure 1A–C**.

**Figure 1.**
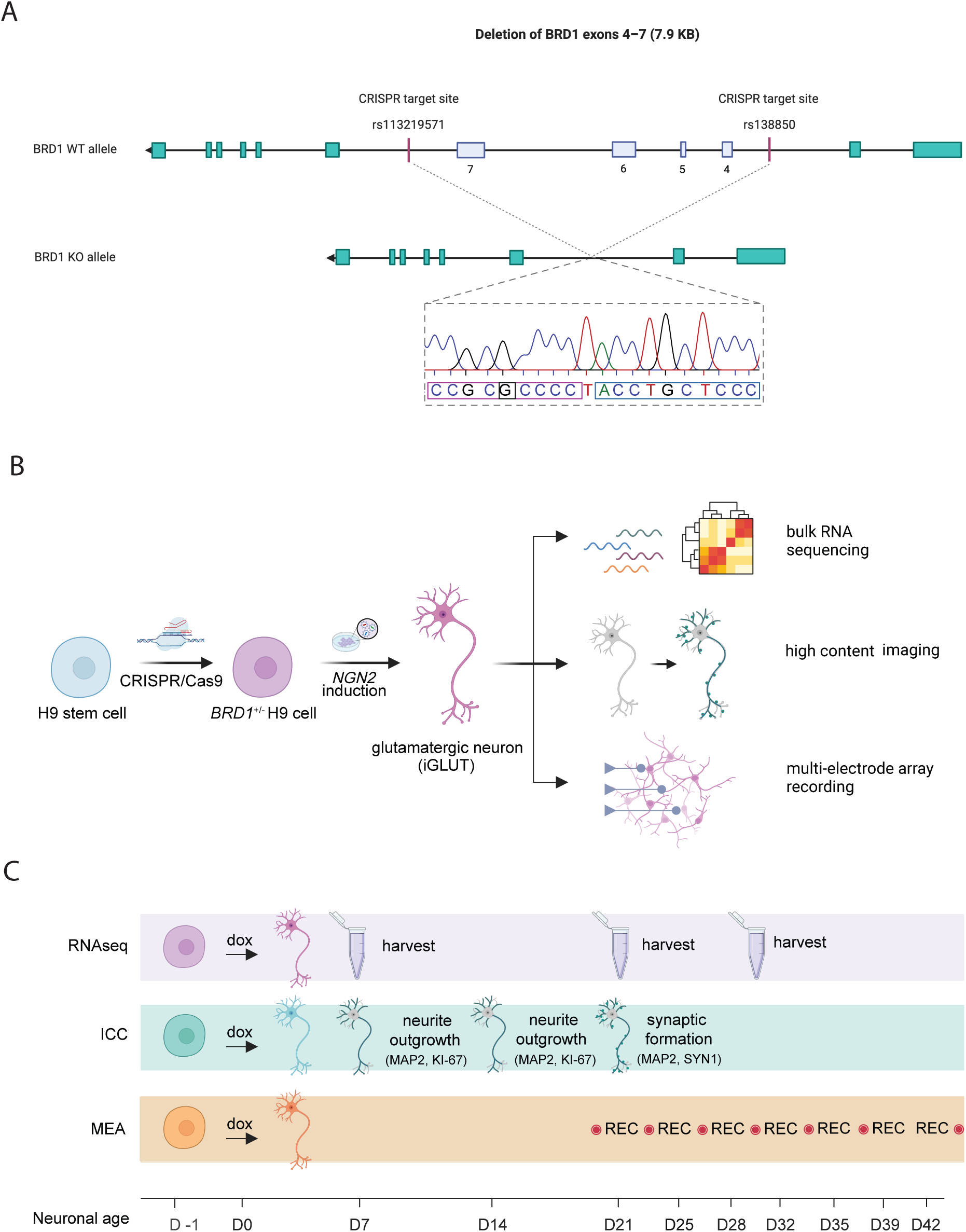
Generation of *BRD1^+/-^* hPSC line and study design. **A.** Validation of mono-allelic deletion of *BRD1* exons 4–7 in H9 hPSCs by Sanger sequencing. With CRISPR/Cas9, mono-allelic targeting of *BRD1* was performed using two sgRNAs targeting individual alle-specific SNP variants in introns flanking exons 4–7. Sequencing data in the purple and blue squares, respectively, match the upstream (SNP rs113219571) and downstream (SNP rs138850) CRISPR target sites, validating the 7.9 KB planned deletion. The highlighted base, G, confirms that the deletion between the two target sites is specific for the allele carrying a G in this position. Likewise, sequencing data from two PCR WT bands (not shown) furthermore confirmed consistent allele-specificity. **B.** Study design. H9 hPSCs were genetically modified using CRISPR/Cas9 to express only one *BRD1* allele, converted into glutamatergic neurons via NGN2-induction and functionally assessed by RNA sequencing, high content imaging and multielectrode array. **C.** Timeline of RNAseq, ICC, and MEA experiments on *BRD1^+/-^* and WT iGLUTs. Experiments were performed in three separate rounds over a course of six months, complementing each other with data retrieved at comparable neuronal ages. Culturing, differentiation and transfer procedure of iGLUTs were kept consistent across experiments where possible to ensure the best conditions for data comparison between experiments.

### hPSC CRISPR targeting and validation

To study the disease phenotypes associated with genetic disruption to the *BRD1* gene on neurodevelopment *in vitro*, we used CRISPR-Cas9-mediated genomic editing to introduce a loss-of-function monoallelic mutation in the *BRD1* gene in a commercially available line of hPSCs (H9). Particularly, we introduced an allele-specific 7.9K bp deletion to remove exons 4–7 of the *BRD1* gene, thereby causing a frameshift predicted to induce non-sense mediated decay of the resulting mRNA product. The deletion was validated by PCR and Sanger sequencing (**Table S1–2** and **Figure S1**).

### Design of guide RNAs

Two CRISPR guide RNAs (crRNAs) with complementary sequences to two allele-specific intron SNP sites (rs113219571 and rs138850) flanking exons 4–7 of the *BRD1* gene were designed using the open-access web tool CRISPOR.org (40) to assess off-target scores, guide efficiency and specificity (**Table S1)**.

### Transfection and clonal expansion

Transfection of H9 cells followed by single-cell FACS sorting was performed to generate a heterozygous clone. Specifically, H9 hPSCs were cultured on feeders and maintained in KnockOut Serum Replacement (KSR) media in a 35 mm center-well plates. On Day 8 post passaging, hPSCs were dissociated for 5–10 min at RT in 1 mL PBS^−/−^ with 0.1% EDTA and harvested in 1 mL E8 media with 1% ROCK inhibitor. Cells were counted (Countess) and 50,000 live cells were plated onto a 35 mm dish on laminin overnight. Culture medium was then substituted with E8 without ROCKi and cells transfected with the GFP plasmid and two allele-specific crRNA guides (**Table S1)**, tracrRNA and Cas9 protein (RNP complex) using Lipofectamine Stem Reagent. Cells were incubated for 48 h, with an extra 500 μL of E8 media added after 24 h. Transfection media was replaced by fresh medium, and 72 h post transfection, cells were dissociated in 30 μL Accutase for ∼8 min, harvested in an additional 500 μL E8 media, centrifuged for 5 min at 300×g and resuspended in 400 μL E8+P/S+ROCKi. Using conservative gating criteria, live, GFP-positive cells were single-cell FACS sorted into two laminin-coated 96-well plates with cloning media (E8+10% CloneR2 supplement) and incubated for ten days at 37°C, 5% CO2 to recover and expand.

### Screening of clones for heterozygosity

Ten days post FACS sorting, 39 healthy clones were dissociated in Accutase, washed and replated into 24-well plates and simultaneous PCR screening for heterozygous clones was performed. Genomic DNA from each clone was extracted using GeneJet Genomic DNA purification kit (Cat K0721, Thermo Fisher Scientific), and regions of interest were subsequently amplified by standard HotStart PCR amplification with the following program: initial pre-denaturation (95°C for 15 min); 36 cycles of denaturation (95°C for 20 s), annealing (59°C for 20 s) and extension (72°C for 20 s); post-termination (72°C for 5 min) and hold at 4°C. PCR products were analyzed by gel electrophoresis on a 2% agarose gel at 80V for 40 min. Of 39 clones, two clones showed bands representing a large deletion of the anticipated size. PCR gel bands representing the WT and *BRD1^+/^* alleles of heterozygous clones (D10 and G10) were purified (Qiagen gel purification kit) and validated in terms of clonality and deletion by Eurofins Genomics using Sanger Sequencing (**Table S2**).

### Validation of clonality and heterozygosity

Sanger sequencing validated one clone, G10, as clonal, simultaneously displaying an allele-specific deletion consistent with the anticipated CRISPR activity **(Figure S1B).** The G10 clone was moved to feeders for further maintenance and validation before long term storage in StemCell Banker in liquid nitrogen. Only hPSCs from the G10 clone was used in the experiments of this study.

### Thawing and adaptation of H9 hPSCs

Prior to neuronal conversion, G10 H9 *BRD1^+/−^* and WT hPSCs were cultured and expanded on 1X Geltrex in complete StemFlex media (Gibco catno. A3349301) with 1X Antibiotic-Antimycotic (Gibco catno. 15240062, contains penicillin, streptomycin, and Gibco Amphotericin B). Upon thawing, cell lines were plated on 1X Geltrex in StemFlex media with added ROCK inhibitor, Chroman 1, and carefully adapted over three weeks from feeder-dependent to feeder-free culturing conditions.

### Viral amplification and purification

E. coli glycerol stocks of NGN2, rtTA, MDL, VSVG and REV plasmids were cultured in 5 mL LB medium with ampicillin in bacterial tubes for 18 h at 37°C at 200rpm. Briefly kept at 4°C, viral cultures were then transferred to Erlenmeyer flasks in 180 mL of LB medium with ampicillin and incubated at 37°C with agitation (160rpm) for 18 h. The flasks were then kept on 4°C for 24 h and virus was purified by maxiprep (Invitrogen PureLink Maxiprep Kit ref. no. K210017) according to the manufacturer’s protocol, with isopropanol added to the eluate for 48 h to achieve a higher yield,

### Viral packaging

Purified virus was transfected into 90% confluent 15 cm dishes with 293HEK cells for packaging. A concentration of 3.1 μg REV, 4.1 μg VSVG, 8.1 μg MDL, 12.2 μg NGN2 or rtTA plasmid was used for each 15 cm dish of 293HEK cells and added to 250 μL of OptiMEM at RT. 110 μL of PEI and 2500 μL of Opti-MEM at RT was mixed in a separate tube and the virus was added carefully to the PEI solution one drop at a time to avoid precipitation of virus. The complete transfection solution was then distributed evenly onto each 15 cm dish with HEK cells in 15 mL HEK media and a full media change was performed 5 h post-transfection. HEK media containing packaged virus particles was collected and replaced with new media 48 h and 72 h after the first media change. Next, the virus-containing media was filtered and distributed evenly into 50 mL tubes and concentrated with Lenti-X concentrator at a final concentration of 25% and stored at 4°C overnight. The virus-containing HEK media was then centrifuged for 45 min at 1500 ×g at 4°C, after which the supernatant was removed, and the pellet resuspended in 500 μL pure DMEM.

### Neuronal conversion of H9 stem cells into induced glutamatergic neurons

Third-generation lentiviral NGN2 transduction of preconditioned PSCs to generate glutamatergic neurons (iGLUTs) was initiated once cells were abundant (D-1). PSCs were dissociated in Accutase, counted (Countess), and transduced by adding 30 μL rtTA and 30 μL NGN2 per 1M cells, after which they were plated and cultured in 1X Geltrex-coated 6-well plates (1M cells/well) in Stemflex media with ROCK inhibitor (Chroman 1). One day post transduction (D0) transgenes were activated with doxycycline (2 μg μL−1) in induction media (DMEM/F12 with Glutamax supplement, 1% N-2, 2% B27–RA, 1% Antibiotic-Antimycotic), and successfully transduced neurons were selected for with geneticin (0.5 mg mL−1) in induction media 48 h post transduction for 48 h (D1-D3). See **Figure S2** for successful transfection and induction in iGLUTs D-1–D1. Three days post induction (D3) neurons were counted and replated in new 4X Geltrex-coated plates in neuron media (Brainphys neuron medium, 1% N2, 2% B27–RA, BDNF (1:1000 dil), GDNF (1:1000 dil), cAMP (1:1000 dil), L-ascorbic acid (1:1000), 1mM Sodium Pyruvate, 1% Antibiotic-Antimycotic, Natural Mouse Laminin). Prior to immunostaining and multielectrode array, *BRD1^+/−^* and WT PSCs were transduced with lentivirus in three independent rounds to correct for anticipated variation in cell-to-virus ratio.

### Molecular, structural and functional phenotyping

#### RNA-seq analysis

RNA was extracted from cultures at D7, D21, and D28 using TRIzol, with six biological replicates per condition. RNA-seq libraries were prepared using the NEBNext Single Cell/Low Input RNA

Library Prep Kit and sequenced on an Illumina NovaSeq 6000 platform (paired-end 150 bp reads, ∼40 million reads per sample) by the Yale Center for Genome Analysis (YCGA).

Quality control on the raw fastq files was performed using fastQC (41). The raw sequences were trimmed using AdapterRemoval (2.3.3). Following standard practices, reads were mapped to human reference genome hg38 (42) using STAR aligner (2.7.11) (43) with --sjdbOverhang set to 150 and default parameters for the rest. Transcript quantification was performed with featureCounts (v2.0.6).

Differential gene expression was calculated in the R programming environment using DESeq2 (v1.40.0) (44) where genes with 0 or 1 total counts for all samples where prefiltered and were excluded from the analysis. Gene-set enrichment analysis (GSEA) based on GO, KEGG and Reactome gene-sets were done using ClusterProfiler (v4.12.6) (45). For enrichment of disease-related gene sets, published psychiatric disorder genetic and genomic disease-related gene lists for PTSD, MDD, schizophrenia (SCZ), BD, ASD, epilepsy and intellectual disability (ID) were used (19). All P values from all gene sets were adjusted for multiple testing using the Benjamini–Hochberg procedure, using a P value< 0.05 threshold to determine significance.

Enrichment analysis of Transcription factor binding sites (TFBSs) enrichment analysis was carried out on DEGs according to Gearing et al.[93] using CiiiDER. Briefly, promotor sequences (2000 bp upstream of TSS) were extracted from the Homo sapiens GRCh38.94 genome file. Identification of TFBSs in these sequences was performed with HOCOMOCOv11_core_HUMAN transcription factor position frequency matrices (downloaded from the HOCOMOCO collection) (46) and a deficit cut-off of 0.15. CiiiDER enrichment analysis of overrepresented DEG TFBSs in DEG query sequences compared to non-DEG query sequences (from 1000 genes with p∼1 and logFC∼0) was determined by comparing the number of sequences with predicted TFBSs to the number of those without, using a Fisher’s exact test.

Cell-type deconvolution/digital cytometry was performed using CIBERSORT tool (65) and single-cell RNAseq reference datasets from neural precursor cells from day 0, 3, 7 and 14 (47) to estimate shifts in cellular composition.

### Immunostaining and high-content imaging microscopy

#### Neurite Outgrowth and Synaptic Assays

Immature D3 *BRD1^+/−^* and WT iGLUTs were seeded onto 4X Geltrex-coated 96-well plates and maintained in neuron media containing Chroman I (1:10,000), doxycycline (1 µg/mL), AraC (4 µM), and geneticin (0.5 mg/mL). For the neurite outgrowth assay, 15K cells/well were plated directly onto Geltrex; for the synaptic assay, 150K cells/well were plated onto a layer of primary human astrocytes (12K cells/well) with 10% fetal bovine serum added to the media. A full media change was performed on D4 to withdraw geneticin, followed by full media changes every 2–3 days until D10 and half media changes thereafter until fixation.

For both assays, cultures were washed with PBS^+/+^ (with Ca^2+^ and Mg^2+^) and fixed in 4% paraformaldehyde/sucrose in PBS^+/+^ for 10 min at RT (neurite assay: D7; synaptic assay: D21).

Fixed cells were washed twice in PBS and permeabilized/blocked in PBS^+/+^ containing 2% normal donkey serum (NDS) and 0.1% Triton X in the dark for 2 h at RT.

Primary antibody incubation was performed overnight at 4 °C in PBS^+/+^ with 2% NDS. MAP2/ck (1:5000; Abcam ab5392) was used for both assays; KI67/rb (1:500; Abcam ab15580) was included for the neurite assay, and Synapsin-1/ms (1:500; Synaptic Systems #106011) for the synaptic assay. Followed by three PBS^+/+^ washes, cultures were blocked for 1 h in PBS^+/+^ with 2% NDS, incubated for 1 h at RT and washed again in PBS^+/+^ before they were incubated with secondary antibodies (1:500 donkey anti-chicken Alexa 647, Thermo Fisher Scientific, A78952 and 1:500 donkey anti-rabbit Cy3, JacksonImmunoResearch, art. no. 711-165-152 for neurite assay; 1:500 donkey anti-chicken, Alexa 647, Thermo Fisher Scientific, A78952 and 1:500 donkey anti-mouse Cy3, Jackson 715-165-151 for synaptic assay) in PBS^+/+^ with 2% NDS for 1 hour at RT. Finally, cultures were washed 3x with PBS^+/+^, with a slightly longer second wash containing 1 μg/ml DAPI and 2% NDS. Imaging for both assays was performed on a CellInsight CX7 HCS Platform using a 20x objective (0.4 NA). Neurite morphology was quantified using the neurite tracing module in Thermo Scientific HCS Studio 4.0 Cell Analysis Software, whereas synapse formation was quantified using the synaptogenesis module.

#### Replicates and statistical analysis

In both assays three different rounds of neuronal conversion were performed for each cell line using different colonies for each new round, serving as biological repeats of the experiment. In the neurite outgrowth and synaptic assay, a total of 20 and 15 identical replicates, respectively, distributed on two 96-well plates and representing each conversion performed (**Figure S3**), were imaged. Nine images were acquired per well for neurite tracing or synaptic analysis, from which either a well average or well sum was calculated per well to represent each replicate. All replicate values within the two genotypes were then listed as equal replicates in a column analysis in GraphPad Prism. For each variable compared, an unpaired, parametric t-test assuming Gaussian distribution was performed on the respective column analysis, followed by a Welch’s correction if variances differed significantly. In cases where data from one or both genotypes did not follow Gaussian distribution, a nonparametric Mann-Whitney (MW) test was performed.

### Multielectrode array

One day before replating D3 iGLUTs, primary human astrocytes (pHAs, Sciencell, no.1800; isolated from fetal female brain) were seeded at 12K cells/well on 4X Geltrex-coated 48-well MEA plates (catalog no. M768-tMEA-48W; Axion Biosystems) in astrocyte medium. The day after, immature D3 *BRD1^+/−^* and iGLUTs were seeded on the astrocytes (150K cells/well) and cultured in neuronal media supplemented with 10% fetal bovine serum. Full media changes were performed every two–three days until D10, after which half media changes were performed twice a week until D46. From D10, AraC and doxycycline was gradually withdrawn by half media changes moving forward. Starting at D21, the electrical activity of iGLUTs was recorded at 37°C for 5 min twice every week using the Axion Maestro MEA reader (Axion Biosystems) (**Figure 1C**), with media changes consistently performed the day before recordings. Using the summary csv-files generated by the Axion Biosystems software, we compared the well averages of “number of spikes” and “number of bursts” per 5 min recording between the two genotypes. For plate layout, see **Figure S3**.

## Results

### BRD1 haploinsufficiency affects neural progenitor transition and early neuronal development

To investigate the impact of BRD1 haploinsufficiency on neuronal development, we introduced a mono-allelic disruptive mutation in hPSCs (**Figure 1A and 2A**), validated targeted on-site CRISPR editing of one clonal hPSC line by Sanger sequencing, and converted isogenic *BRD1^+/-^* and wildtype hPSCs, matched by passage number, into induced glutamatergic neurons (iGLUTs) using *NGN2* overexpression (36) (see **Figure 1B-C** for study design). As early as D7, MAP2-positive cells with neuronal morphological features were evident in both genotypes (**Figure 2E**). Analysis of *BRD1* expression in WT iGLUTs revealed a peak at D7 followed by a gradual decline towards D28 (t-test, p_D7-D28_ = 0.03). In contrast, and consistent with monoallelic disruption, *BRD1* expression was significantly reduced in *BRD1*^+/-^ iGLUTs (**Figure 2A**, 2-way ANOVA, p_genotype_=0.006), with no apparent temporal regulation of its expression.

**Figure 2.**
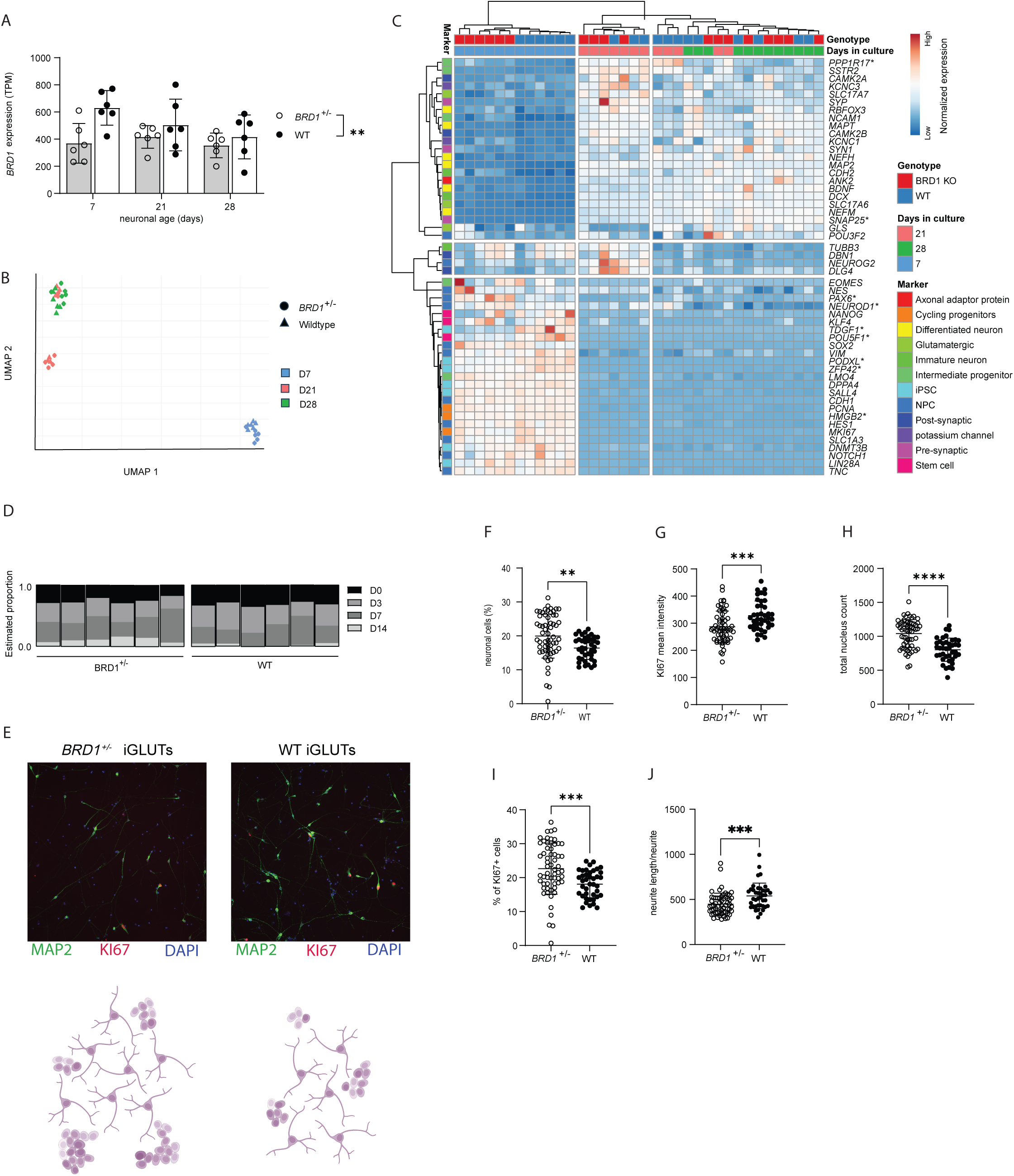
BRD1 haploinsufficiency affects neural progenitor transition and early neuronal development. **A.** Normalized expression of BRD1 in *BRD1^+/-^* and WT iGLUTs during neuronal development. Error bars indicate mean ± SD. **B.** UMAP trajectory plot, showing cell state transitions of NGN2-induced iGLUTs from D7 to D28. **C.** Heatmap of normalized expression of curated cell state markers in WT and *BRD1^+/-^* iGLUTs across D7, D21 and D28. Asterisks indicate markers with significant genotype by time interaction after FDR correction (two-way ANOVA). **D.** *In silico* deconvolution analysis of *BRD1^+/^* iGLUTs, using single-cell RNAseq reference data from neural precursor cells. **E.** Comparison of *BRD1^+/-^* and WT iGLUTs at D7 (neurite outgrowth). Top panel: MAP2 and KI67; lower panel: schematic of iGLUT morphology. **F–J.** Neurite outgrowth assay (ICC) on D7 iGLUTs. All data points represent the mean of nine snapshots per well. Error bars indicate mean ± SD. **F.** Percentage of mature neurons (MAP2-positive cells) of total cell count. **G.** Mean intensity of proliferation marker KI67 per KI67-positive cell. **H.** Total nucleus count (well sum). **I.** Percentage of KI67-positive cells of total cell count. **J.** Total neurite length per neurite.

Transcriptomic profiling at D7, D21 and D28 revealed tight clustering of biological replicates within each time point (**Figure 2B**). At D7, coinciding with the peak of *BRD1* expression in WT cells and the largest relative difference between genotypes, *BRD1*^+/-^ and WT iGLUTs segregated into distinct clusters (**Figure S4**). This early divergence indicates that BRD1 haploinsufficiency perturbs the transcriptional program that normally coordinates the initial stages of neuronal differentiation.

To examine whether early differences reflected altered developmental trajectories, we assessed the expression dynamics of curated cell state markers across the three time points (**Figure 2C**). Several pluripotency- or early progenitor-associated genes (*POU5F1*, *ZFP42*, *TDGF1, PODXL, HMGB2*) and markers linked to early neuronal commitment or maturation (*PAX6*, *NEUROD1*, *PPP1R17*, *SNAP25*) showed significant genotype-by-time interactions after FDR correction (2-way ANOVA, p_FDR_ < 0.05; **Table S3**), indicating an accelerated transition toward neuronal differentiation in *BRD1*^+/-^ cultures. This interpretation was supported by *in silico* deconvolution results, which indicated that *BRD1^⁺/⁻^* iGLUTs at D7 were more closely aligned with transcriptional states characteristic of more advanced neuronal maturation compared with their WT counterparts (**Figure 2D**, p_D14_**=**0.0505).

Consistent with accelerated transition, high content imaging based immunocytochemical (ICC) analysis at D7 showed a higher proportion of MAP2-positive cells (**Figure 2E and F**, MW test, p=0.001) and reduced Ki67 intensity in *BRD1*^+/-^ iGLUTs compared to WT (**Figure 2E and G**, t-test, p=0.0005). *BRD1*^+/-^ iGLUTs also exhibited a greater total nucleus count (**Figure 2H**, t-test, p<0.0001) and a higher proportion of Ki67-positive cells at D7 **(Figure 2E and I**, Welch’s t-test, p<0.0002). Neurite length was significantly reduced in *BRD1*^+/-^ iGLUTs compared to WT (**Figure 2E and J**, MW test, p=0.0003), independent of neurite or branch point number **(Figure S5**).

Collectively, these findings point to an aberrant neurodevelopmental trajectory in *BRD1*^+/-^ iGLUTs, marked by increased progenitor proliferation alongside early expression of neuronal markers, suggesting a decoupling of proliferation control and differentiation timing, resulting in disrupted neurodevelopment.

### BRD1 deficiency disrupts transcriptional control of neuronal maturation and impacts neurodevelopmental disorder-associated gene networks

Transcriptomic analysis revealed 373 differentially expressed genes (DEGs) in *BRD1*^+/-^ iGLUTs compared to WT across development (**Table S4**), with the most pronounced differences seen at D7 (**Figure 3A** and **Table S5**). 18 DEGs showed consistent directional changes at all three timepoints (D7, D21 and D28) (**Figure 3B** and **Table S5–7**), including several genes previously implicated in neurodevelopmental disorders, such as *MAGEE1* and *MIR325HG* (ASD candidates) (48)(41) and *GRIA3*, which has been linked to intellectual disability (ID), schizophrenia (SZ), and ASD **(Figure 3C)** (49–52). Other consistently dysregulated genes included the stress-related corticotropin-releasing factor-encoding gene, *CHR*, transcription factors such as HESX1 and TCEAL5, and multiple zinc finger proteins (ZNF37A, ZNF585A, ZNF248, ZNF959), suggesting widespread disturbance of gene regulatory networks. Further connecting transcriptomic changes to the observed neurodevelopmental phenotype, promoter sequences of identified DEGs were significantly enriched (p_val_<0.05) for consensus transcription factor binding sites (TFBSs) of key regulators of neural differentiation, including POU5F1, POU3F2, KLF4, and ZFP42 (**Figure 3D** and **Table S8**). Notably, we also observed enrichment for TFBS of several nuclear receptors previously shown to interact with BRD1 (53), including PPARG, ESR1 and NR3C2 (**Figure 3D**). Whereas no significant gene-set association to neuropsychiatric disorders were found for DEGs using the most recent GWAS data for SZ, ASD, BPD, MDD and ADHD, gene set enrichment analysis (GSEA) using disease-related gene lists revealed psychiatric disorder enrichment for PTSD, MDD, SZ, BD, ASD, epilepsy and intellectual disability (ID) (**Figure 3E** and **Table S4**) (54).

**Figure 3.**
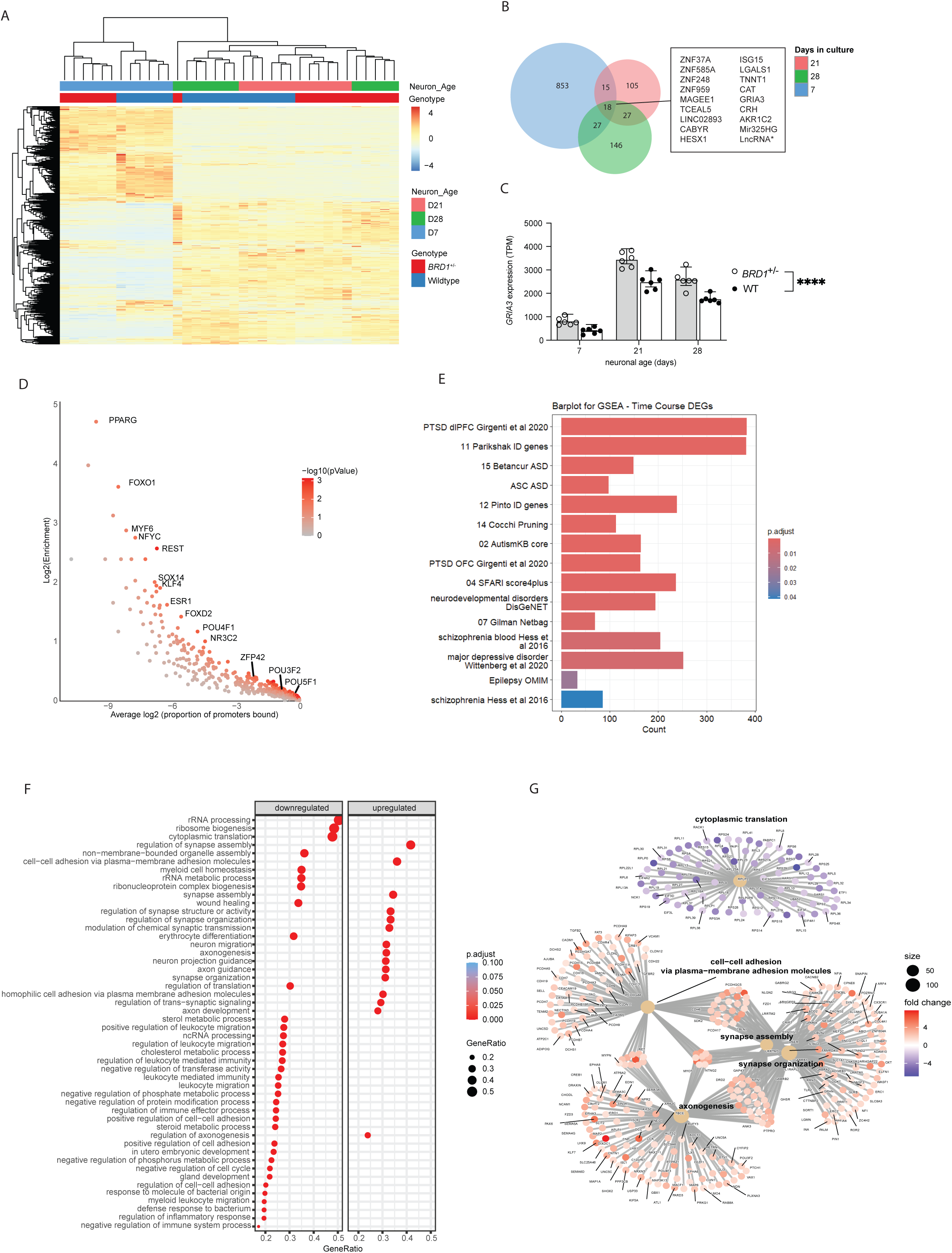
BRD1 deficiency disrupts transcriptional control of neuronal maturation and impacts neurodevelopmental disorder-associated gene networks. **A.** Heatmap showing DEGs (p_adj_<0.05) between *BRD1^+/-^* and WT iGLUTs at D7, D21 and D28. **B.** Venn diagram illustrating the overlap of DEGs at D7, D21 and D28. **C.** Normalized expression of *GRIA3* in *BRD1^+/-^* and WT iGLUTs during neuronal development. Error bars indicate mean ± SD. **D.** CiiiDER enrichment analysis of transcription factor binding sites among DEGs. **E.** GSEA of genes with LoF mutations in neuropsychiatric disorders. **F.** GSEA highlighting associated biological processes (BPs) of DEGs in *BRD1^+/-^* iGLUTs. **G.** Gene network visualization of BPs from GSEA shown in E. BPs are grouped into gene networks based on functional similarity and shared genes, illustrating their interconnectivity. Node size reflects the number of genes in each BP, while the colors purple and red represent downregulation and upregulation, respectively.

GSEA performed to characterize the broader biological processes affected by BRD1 haploinsufficiency, revealed consistent enrichment of gene networks involved in neuronal development and function (**Figure 3F**), including synapse assembly and organization, cytoplasmic translation, as well as axonogenesis and cell-cell adhesion **(Figure 3G)**. Similar pathway enrichment patterns were observed when GSEA was performed for each time point independently (**Figure S6A–C** and **Table S5–7**), underscoring the sustained disruption of key neurodevelopmental processes in in *BRD1^+/-^* iGLUTs.

#### BRD1 deficiency alters structure, composition and activity of mature neuronal networks

Cell composition and morphological characteristics of mature neurons at D21 was assessed by high content imaging. Indications of accelerated neurogenesis at D7 (**Figure 2**) were no longer apparent by D21. Instead, *BRD1*^+/-^ iGLUTs displayed significantly lower proportion of MAP2-positive neurons compared to WT (**Figure 4A** and **C**, t-test, p=0.025), along with fewer neurites per neuron (**Figure 4A** and **D**, MW test, p<0.0001). However, total neurite length (**Figure 4A** and **E**) and branching complexity **Figure 4A** and **F**) were not significantly different between genotypes. In accordance with the broader dysregulation of network of genes implicated in synapse assembly and organization, *BRD1* haploinsufficiency resulted in synaptic puncta that were significantly smaller (**Figure 4A** and **G**), although synaptic density remained unchanged in *BRD1*^+/-^ iGLUTs (**Figure 4A** and **H**). To assess functional maturation, we used multielectrode array recordings to measure neuronal activity across development. Although immature neurons (D21) showed similar neuronal activity, a significant increase in *BRD1*^+/-^ iGLUT spike activity (**Figure 4I)** (but not bursts **Figure 4J**) was evident with extended maturation (D32-D42) (2-way RM ANOVA, p=0.0349). Together, these findings suggest that BRD1 haploinsufficiency leads to reduced structural complexity, altered synapse morphology, and dysregulated neuronal activity in maturing networks.

**Figure 4.**
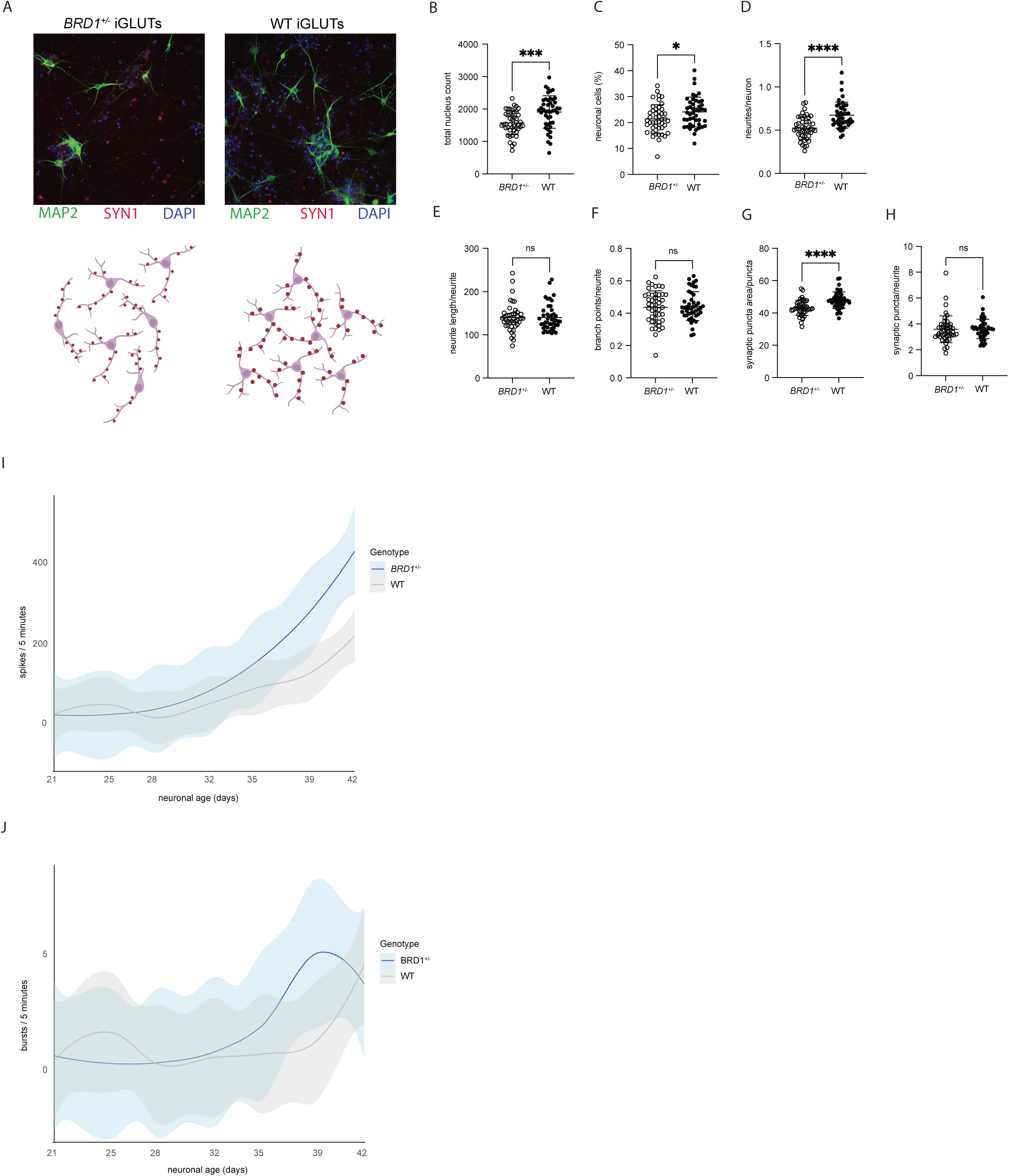
BRD1 deficiency alters structure, composition and activity of mature neuronal networks. **A.** Comparison of *BRD1^+/-^* and WT iGLUTs at D21 (synaptic formation). Top panel: MAP2 and SYN1; lower panel: schematic of iGLUT morphology. **B–H.** Synaptic formation assay (ICC) on D21 iGLUTs. All data points represent the mean of nine snapshots per well. Error bars indicate mean ± SD**. B.** Total nucleus count (well sum). **C.** Percentage of mature neurons (MAP2-positive cells) of total cell count. **D.** Total number of neurites per neuron. **E.** Total neurite length per neurite. **F.** Total number of branch points per neurite. **G.** Synaptic puncta area per puncta. **H.** Total number of synaptic puncta per neurite. **I–J.** MEA spike and burst rates over time. Rates are plotted as averages per well of total spike/burst number per well recorded from 16 electrodes over 5 min. **I.** LOESS plot showing MEA spike rates in iGLUTs from D21–D42. **J.** LOESS plot showing MEA burst rates in iGLUTs from D21–D42.

## Discussion

Disruptions in neurodevelopmental processes contribute to the etiology of major psychiatric and neurodevelopmental disorders. BRD1 has emerged as a key transcriptional regulator implicated in these conditions, yet its functional role in pathophysiological processes remains unclear. In this study, we investigate the impact of BRD1 haploinsufficiency on neuronal differentiation and maturation using human glutamatergic neurons (iGLUTs). *BRD1*^+/-^ neurons exhibit asynchronous neurodevelopment, with an accelerated transition from progenitor to neuronal states followed by impaired neuronal network complexity and reduced neuron density in more mature cultures. Transcriptomic analysis reveals persistent dysregulation of genes associated with synapse organization, and NDDs, including SZ, ASD, and intellectual disability. Despite this, BRD1 haploinsufficiency did not impact density of synapses, but left a functional impact as evident from altered electrophysiological activity patterns in late stages of culturing. These findings highlight a critical role for BRD1 in orchestrating early neuronal programming and maintaining structural and functional integrity in developing neural circuits. Combined with previous evidence of BRD1’s role in embryonic stem cell differentiation through its function as a scaffold in histone acetyltransferase complexes (55), our results extend its importance to human neuronal lineage progression and offer new insights into how BRD1 dysfunction may contribute to the developmental origins of neuropsychiatric disorders.

The importance of temporal and spatial regulation in CNS development is well established. As neural progenitors mature, their ability to generate specific neuronal populations changes over time, involving dynamic changes in gene expression and cellular identity (56–58). Dysregulation in these processes can result in either a premature exit from the progenitor pool or abnormal differentiation trajectories, both of which can contribute to NDDs (59–61). Intriguingly, our observations of over-proliferation of progenitors and accelerated neuronal maturation in BRD1-deficient neurons are strikingly similar to those reported in models of NDDs based on other epigenetic modifiers (15,62–72). For instance, genetic disruption to the ASD-associated gene *CHD8*, which encodes a chromatin remodeler, has also been shown to result in over-proliferation of progenitor cell populations (73), and cause premature differentiation of neuronal progenitors. Similarly, genetic disruption to *SUV420H1*, a chromatin-modifying enzyme that interacts with BRD1, is associated with accelerated maturation of deep-layer projection neurons in an organoid model (15). Collectively, these findings underscore a shared molecular mechanism between chromatin remodelers centered on an underlying dysregulation in the temporal control of neuronal differentiation and contribute to a growing body of literature implicating chromatin remodeling and transcriptional regulation as central mechanisms underlying NDDs (74).

A reduction in neuronal density alongside stable neurite length and branching suggests that *BRD1*^+/-^ iGLUTs may experience deficits in survival or maintenance rather than neuritogenesis per se. This phenotype is reminiscent of findings in models of chromatinopathies such as *KMT2A* and *KDM5C* mutations, where disruptions in chromatin remodeling lead to selective neuronal vulnerability and loss of network complexity over time(75). Similarly, in a mouse model of Brd1 haploinsufficiency, *Brd1*^+/−^ mice (37), structural changes such as a reduction in neuronal density (37), brain volumetric changes (35) and loss of cortical subpopulations of specific interneurons (e.g., parvalbumin-expressing fast-spiking interneurons) have been observed (37). Furthermore, structural alterations in the *Brd1*^+/−^ mouse striatum, particularly a reduction in the number of medium-sized spiny neurons further underscores the notion of selective neuronal vulnerability (35). These changes are consistent with a loss of network complexity and function, as observed in *BRD1*^+/-^ iGLUTs, where reduced neuronal density in the mature cultures may point to a failure to sustain proper transcriptional programs that support neuronal maintenance, potentially leading to a shift in network composition rather than outright structural atrophy.

While synaptic density remained unchanged in *BRD1*^+/-^ iGLUTs, this does not necessarily indicate normal synaptic function or network excitability. Synaptic density alone does not dictate network activity; rather, functional changes such as altered receptor composition, synaptic strength, or neurotransmitter release dynamics can profoundly influence neuronal excitability. Notably, we observed that synapses in *BRD1*^+/-^ iGLUTs were significantly smaller, suggesting potential deficits in synaptic maturation or stability. At the molecular level, we found a significant and persistent upregulation of key genes involved in synapse assembly and organization, including *GRIA3*, which encodes the GluA3 subunit of AMPA receptors, a central mediator of excitatory synaptic transmission. It is thus conceivable that while transcriptional programs supporting synaptic formation are activated, *BRD1*^+/-^ iGLUTs may exhibit qualitative rather than quantitative changes in synaptic connectivity. Alterations in synapse size and receptor composition could shift the balance of excitatory signaling, impacting network function. Such findings are in line with data from *Brd1*^+/-^ mice, where the structural integrity of synapses appeared compromised, as evidenced by alterations in dendritic spine morphology, such as a decrease in the number of mature mushroom-type spines (34)—leading to imbalanced excitation-inhibition, increased seizure susceptibility and reduced inhibitory postsynaptic current frequency (37). The increased neuronal activity in *BRD1*^+/-^iGLUTs observed at later stages of culturing follows a pattern that has been described in other ASD models (76), and could reflect compensatory or maladaptive plasticity mechanisms, where enhanced excitatory transmission partially offsets early deficits in network maturation.

This study establishes BRD1 as a critical modulator of neuronal development, particularly in balancing progenitor proliferation and neurogenic maturation. Our findings reinforce the central role of chromatin remodeling in orchestrating neurodevelopmental timing, highlighting how disruptions in epigenetic regulation contribute to the pathophysiology of NDDs. The convergence of our results with other models of chromatinopathies suggests that impaired temporal control of neuronal differentiation may be a shared mechanism underlying these disorders. Moving forward, further investigation into the molecular interactions between BRD1 and other chromatin modifiers, as well as their downstream effects on gene regulatory networks, will be crucial for elucidating how epigenetic mechanisms shape neurodevelopment. These insights could ultimately pave the way for therapeutic strategies targeting epigenetic regulators to correct abnormal developmental trajectories in NDDs.

## Acknowledgments

This study was supported by research grants from the A. P. Møller Foundation, Helga and Peter Kornings Fond, The Jascha Foundation, Augustinus Foundation, William Demant Foundation, A. P. Møller Foundation, Knud Højgaard Foundation and the Graduate School of Health at Aarhus University. Research funding was also provided by R01MH123155 and RM1MH132648.

We extend our special thanks to Susanne Hvolbøl Buchholdt Seldrup from the Denham Lab for providing indispensable technical and practical assistance regarding cell culture transfection, maintenance and long-term preservation of cells in the generation of our genetically modified cell model. We also thank Sadaf Ghorbani, Olivia Livoti, Ray Rigat and the rest of the Brennand Lab at Yale University.

## Author contributions

Conceptualization: AB, PQ

Methodology: PQ, JD, MD, DP, JEH, PJMD, KJB

Investigation: JD

Visualization: JD, PQ, DPM, JEH

Funding acquisition: JD, PQ, KJB

Project administration: JD, PQ, MD

Supervision: PQ, MD, PJMD, KJB, AB

Writing – original draft: JD, PQ

Writing – review & editing: All authors

## Notes

### Competing Interest Statement

The authors have declared no competing interest.

