## Supplemental figures for "BRD1 haploinsufficiency alters early neuronal programming and disrupts maturation in human induced glutamatergic neurons"

### Supplementary figures

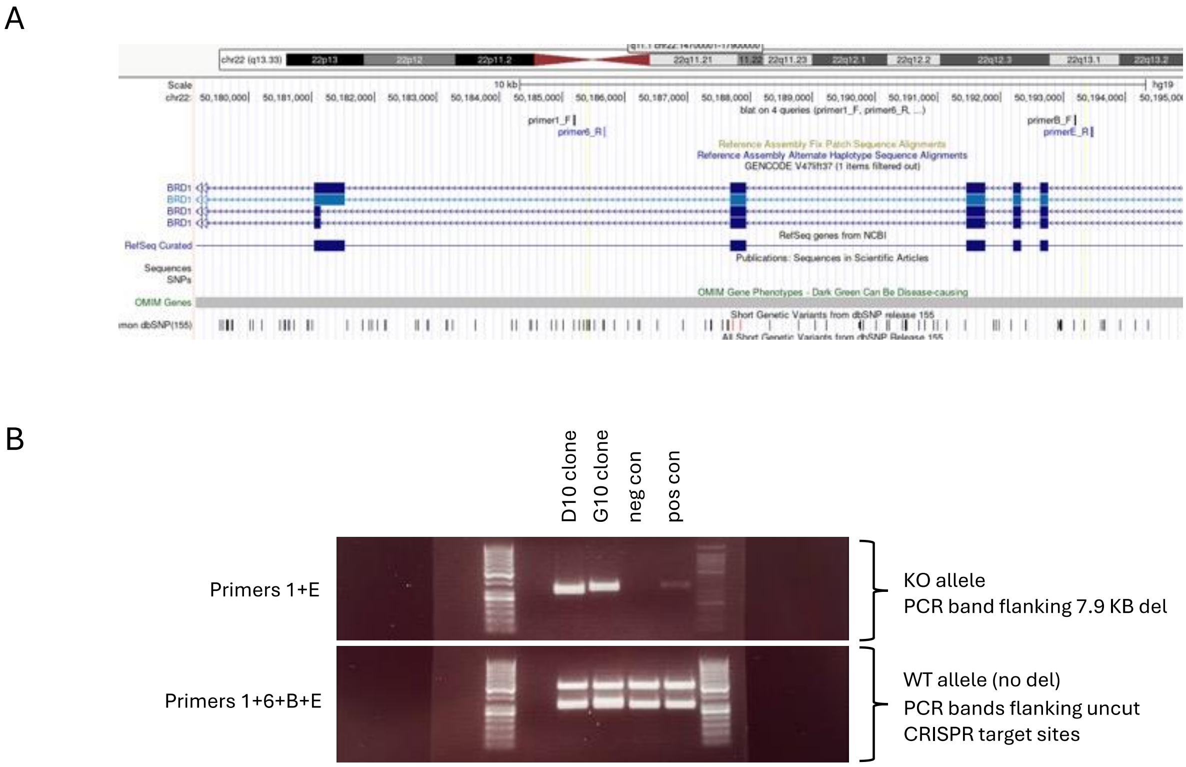


**Figure S1. Validation of monoallelic 7.9 KB deletion of exons 4–7 in *BRD1*. A.** PCR primer positions flanking expected CRISPR cut sites on each side (yellow lines) of the desired 7.9 KB deletion. **B.** PCR validation of deletion. Primer pairs 1+6 and B+E were designed to produce PCR products over the desired cut sites, if no deletion has taken place, thus producing one PCR product per primer pair for the WT allele. Primer pair 1+E were designed to produce a PCR product over the desired deletion, if the deletion has taken place, thus producing one PCR product for the BRD1 deletion allele. On the gel, we see two PCR bands for the WT allele and only one PCR band for the deletion allele of their expected product size, confirming heterozygosity in the clones. Validation of clonality and allele-specificity were confirmed in the G10 clone by Sanger sequencing (Figure 1A), and hPSCs from this clone only was used for all experiments of this study.

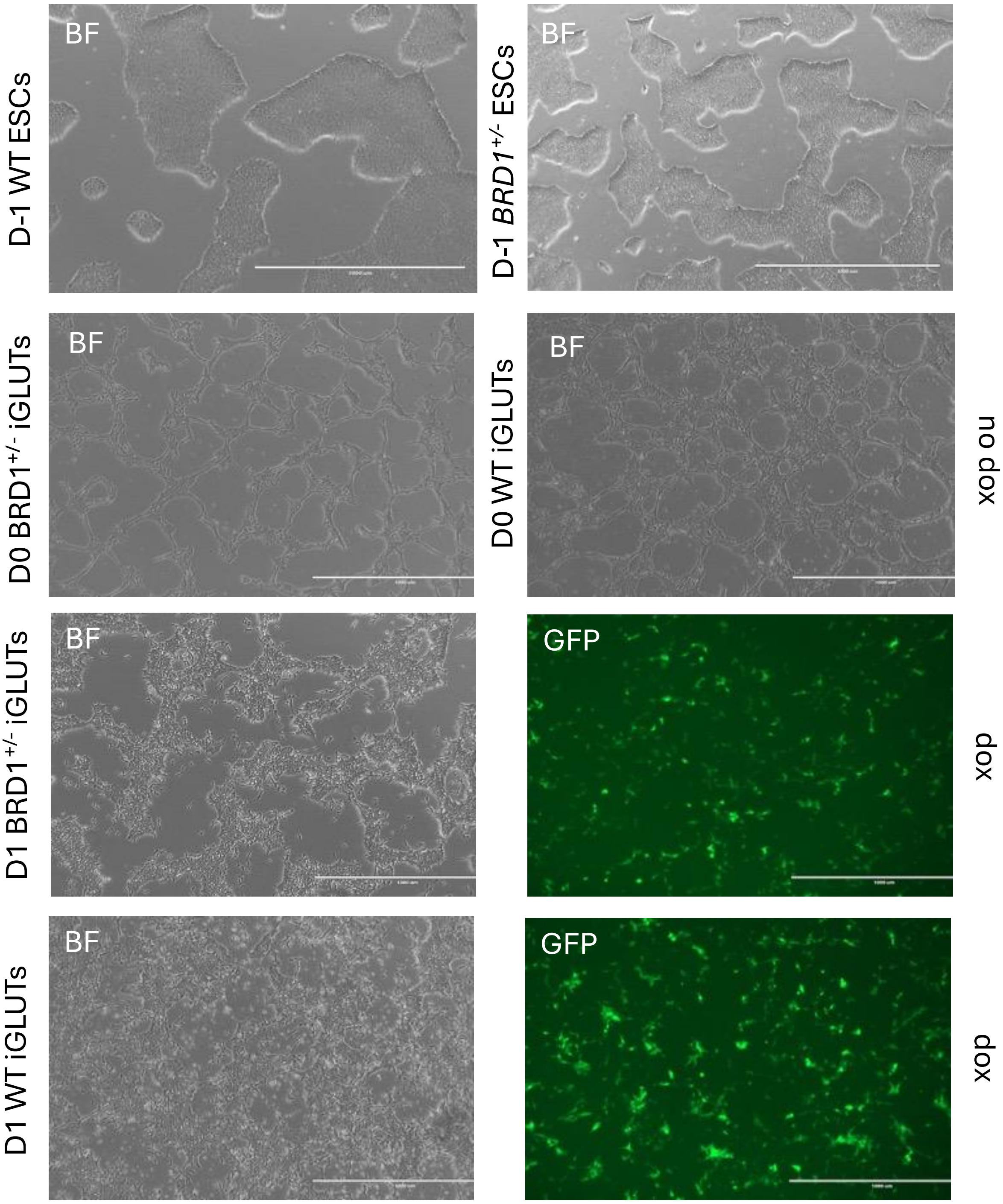


**Figure S2.** Bright-field (BF) and FITC (GFP) images of iGLUTs on day of transduction (D-1), one day post transduction (D0) and one day post dox induction (D1).

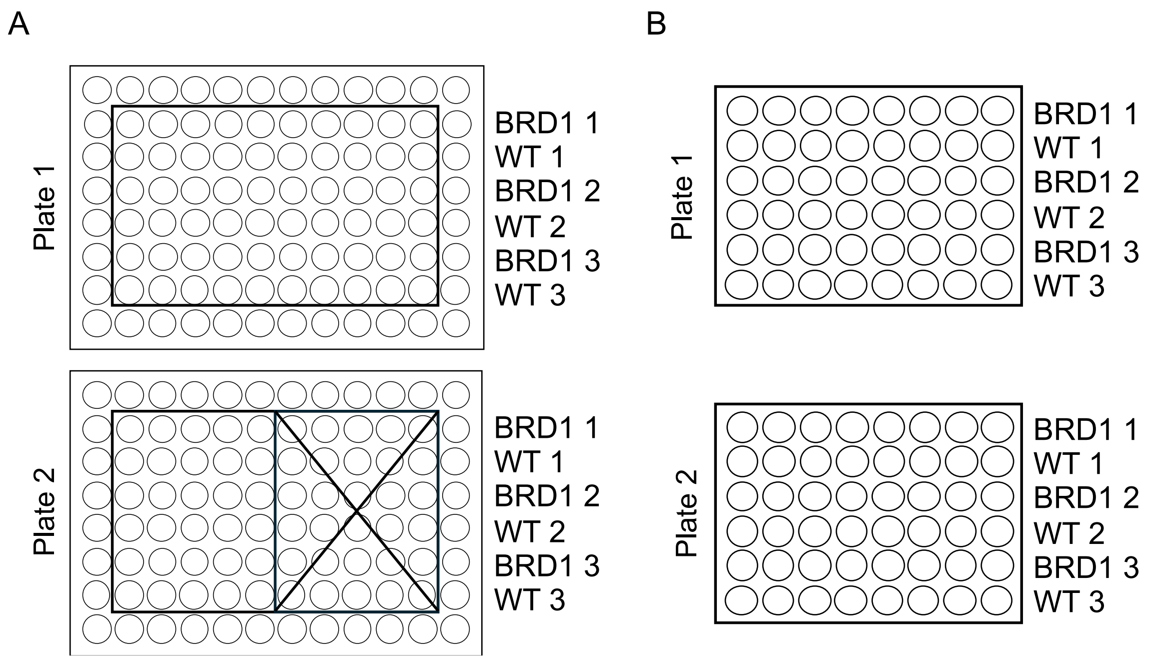


**Figure S3. Plate layout used for ICC and MEA, showing three biological differentiation replicates per genotype.** Human PSCs from the *BRD1*$^{+/-}$ G10 clone was used in all experiments. **A.** ICC plate layout. **B.** MEA plate layout.

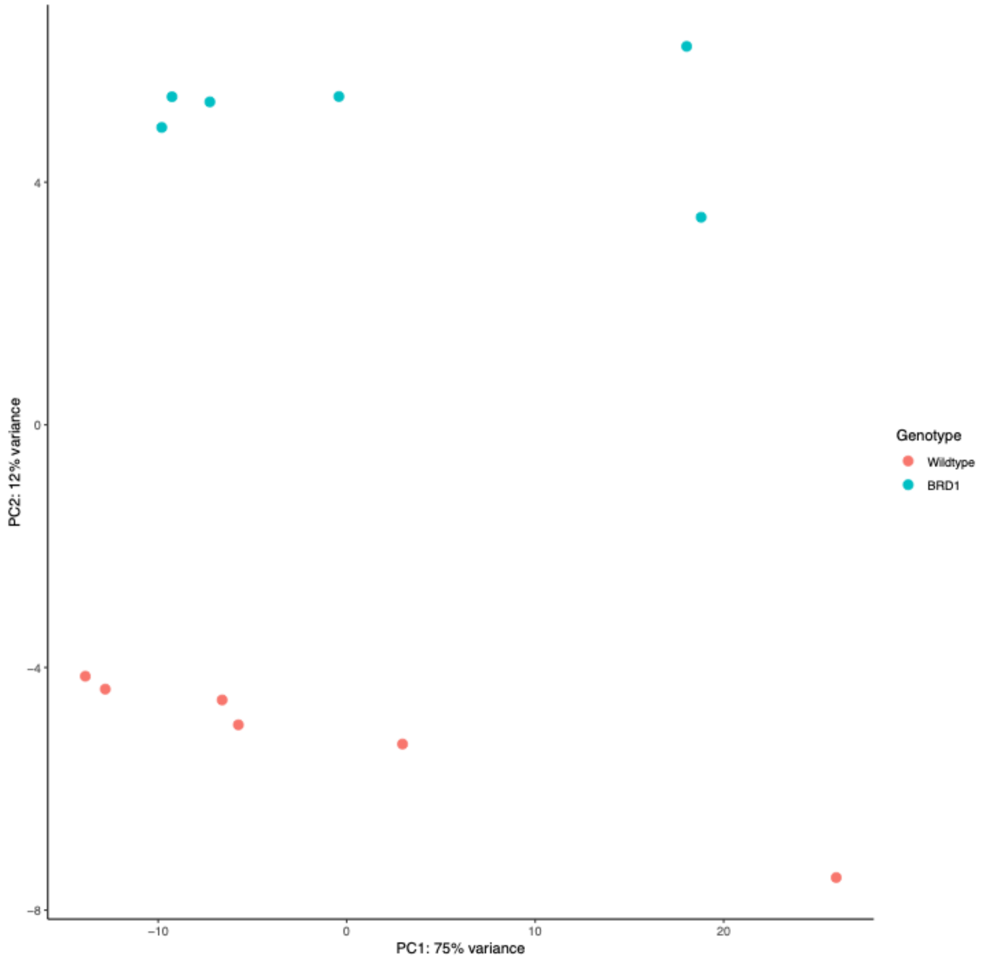

**Figure S4.** PCA plot of D7 iGLUTs.

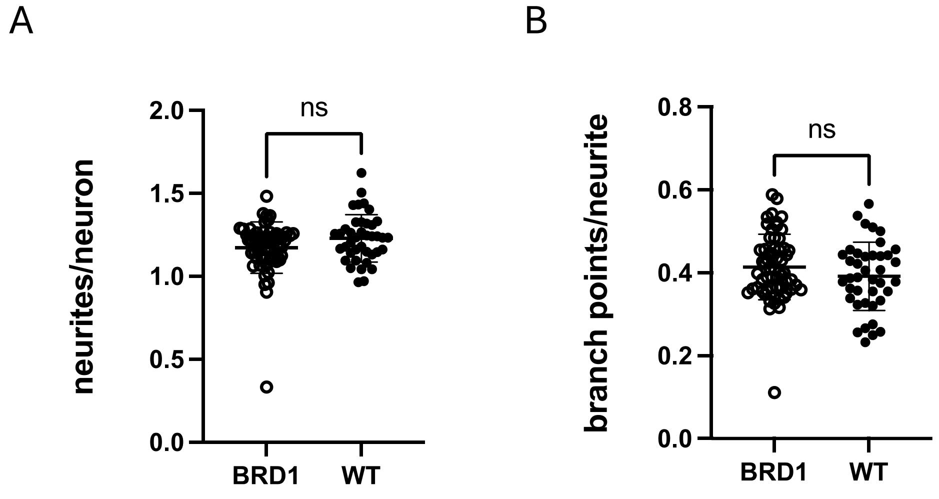


**Figure S5. Neurite outgrowth assay on D7 iGLUTs.** All data points represent the mean of nine snapshots per well. **A.** Total number of neurites per neuron. **B.** Total number of branch points per neurite.

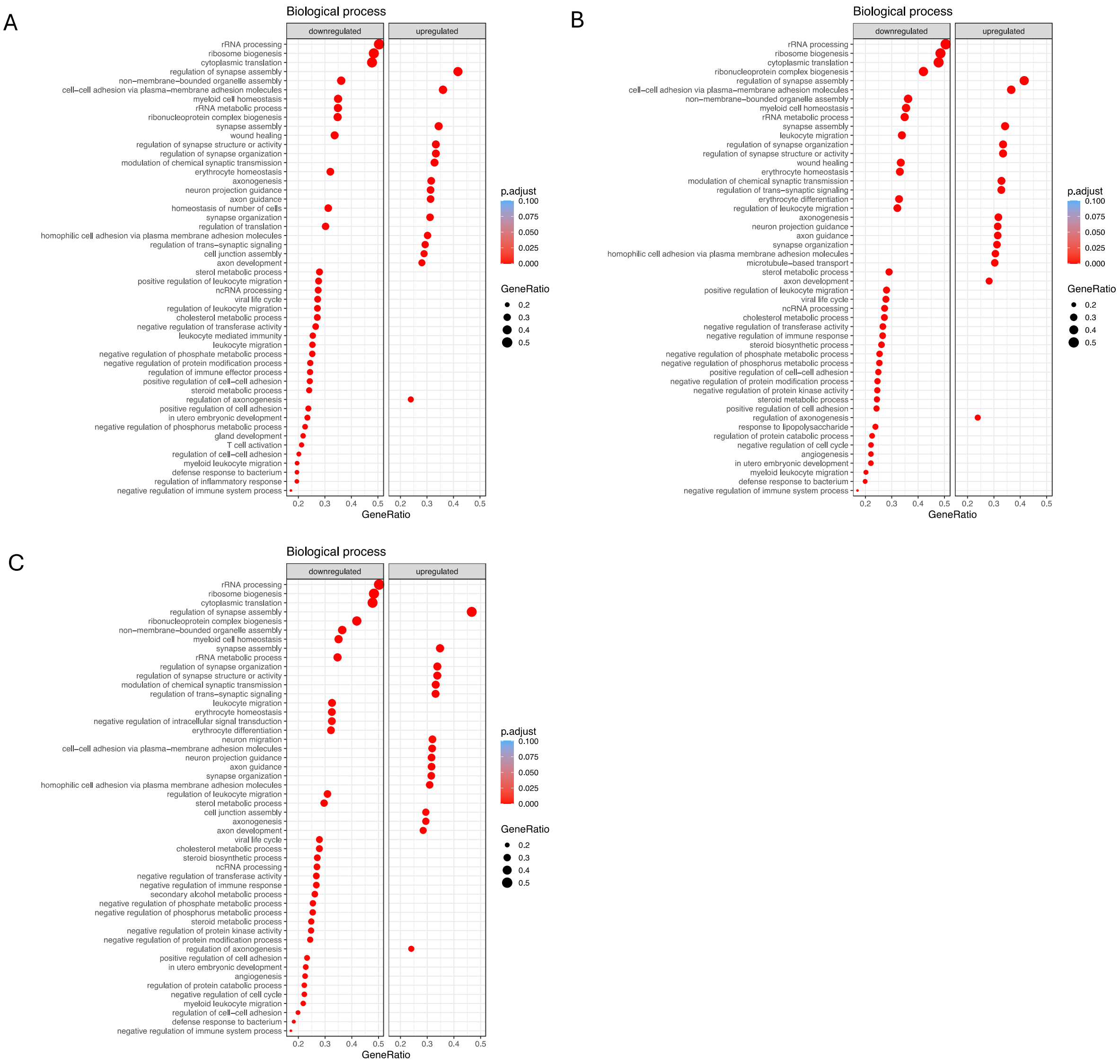


**Figure S6.** GSEA highlighting associated biological processes (BP) of DEGs in *BRD1*$^{+/-}$ iGLUTs at D7, D21 and D28. **A.** GSEA BP of D7 DEGs. **B.** GSEA BP of D7 DEGs. **C.** GSEA BP of D7 DEGs.
